# Effects of a demand optimization intervention on laboratory test utilization in primary care

**DOI:** 10.1101/573956

**Authors:** Magda Bucholc, Maurice O’Kane, Brendan O’Hare, Ciaran Mullan, Paul Cavanagh, Siobhan Ashe

## Abstract

There is evidence of increasing use of laboratory tests with substantial variation between clinical teams which is difficult to justify on clinical grounds. The aim of this project was to assess the effect of a demand optimisation intervention project on laboratory test requesting by general practitioners (GPs) in an area of Northern Ireland supported by the Clinical Chemistry Laboratory service of Western Health and Social Care Trust (WHSCT). The intervention package was developed in conjunction with the Western Local Commissioning Group and consisted of educational initiatives, feedback to 55 individual practices on test request rates with ranking relative to other practices, and a small financial incentive for practices to reflect on their test requesting activity. Overall test utilization rates of profile tests, HbA_1c_, and PSA one year before, during, and one year after the intervention were measured using laboratory databases of the Altnagelvin Area Hospital, Tyrone County Hospital, and the Erne (South West Acute Hospital. The intervention was associated with mixed effects. First, we observed a reduction of 5.1% in the median profile test request rates and a decrease in their between practice variability. The overall downward trend in variability of profile test request rates was found statistically significant (*p* = 0.03). Second, we found a significant increase in both the volume (*p* < 0.0001) and between practice variability (*p* = 0.0001) of HbA_1c_ requests per patient with diabetes. The increase in HbA_1c_ requests may reflect a more appropriate rate of diabetes monitoring and also the adoption of HbA_1c_ as a diagnostic test. Yet, the subsequent 600% increase in between practice variability of HbA_1c_ ordering rates may imply an inconsistent implementation of recommended guidelines by GPs. Finally, there was a 29.3% increase in the median and 35% increase in between practice variability of request rates for PSA, the reasons for which are unclear.

## Introduction

Laboratory testing is an integral part of the clinical decision making process for the disease diagnostics, management, and prognosis [1]. This includes early disease detection, disease surveillance, identification of patients at risk for a disease as well as selection and evaluation of a patient’s treatment based on the results of a lab test [2]. In recent years, the growing use of laboratory tests and in particular, the substantial variation in test ordering rates between clinical teams have become a major concern given rising health care costs [3–4]. The reasons for the increase in the total number of order tests as well as the substantial variation in test ordering between GP practices are still imperfectly understood; however, possible explanations include a lack of knowledge about the appropriate use of individual tests, the use of different clinical guidelines and protocols across GP practices, increased fear of errors and malpractice liability claims as well as professional and practice-related factors, such as GP’s age, GP practice size or type [5–8].

While it is difficult to specify for most laboratory tests what an ‘appropriate’ test request rate might be for a given patient population, it is probable that between-practice variability in test ordering rates is to some extent reflected in inappropriate laboratory utilization through over-requesting (unnecessary repeat requesting of tests), under-requesting (a failure to prescribe clinically indicated testing), and incorrect requesting (selection of an incorrect laboratory test) [9–12]. Several studies showed that around 25-40% of test requests may be unnecessary [13–15], and do not contribute to patient management. While overutilization of laboratory tests drives costs up across the health care system, their under-or incorrect ordering can have serious consequences for the individual patient through failure to diagnose or manage disease optimally [16].

Many attempts have been made to change test ordering performance. A number of studies reported on various clinical interventions designed to improve laboratory utilization and manage demand for laboratory services, with success depending on the medical context, local factors and clinical team engagement [17]. The efforts to improve the appropriateness of laboratory testing behaviour included educational initiatives on the role, limitations, and appropriate retest intervals of individual laboratory tests [18], feedback-based interventions on test usage [19–22], a redesign of laboratory tests request forms [23], and implementation of locally agreed clinical guidelines and electronic medical record prompts for laboratory test orders [24, 25]. A number of studies showed that most of successful strategies for optimizing laboratory demand consisted of a combination of interventions [26, 27].

In this study, we examined whether the volume and variability in laboratory test requests by GPs was reduced by a multifaceted laboratory demand optimisation intervention undertaken as a quality improvement initiative conducted in a primary care setting. In addition, we compared the effects of the intervention on the laboratory test ordering behaviour in GP practices located in rural and urban areas.

## Materials and Methods

### Study setting

The demand optimization intervention was undertaken in 55 separate primary care medical practices within the catchment area of the Northern Ireland (NI) Western Health and Social Care Trust (WHSCT) covering NI local council areas of Londonderry, Limavady, Strabane, Omagh, and Fermanagh. At the commencement of the project, the individual primary care medical practices were composed of between one and eight (mean 3.1) general medical practitioners; eight of the 55 practices comprised a single general medical practitioner. The WHSCT provides laboratory services to these practices with networked laboratories in each of the three large urban centres of Londonderry, Omagh, and Enniskillen. The primary care practices were further situated in either rural or urban areas using data from the Census Office of the Northern Ireland Statistics and Research Agency [24]. Since the NI settlement classification does not give continuous spans of particular area types, a practice was designated as urban if its postal address was situated in a settlement of more than 10,000 residents following the urban-rural classification thresholds used by the Department for Environment, Food and Rural Affairs (DEFRA) and the Department for Communities and Local Government (DCLG) [28]. Under this definition, 31 practices were designated as urban and 24 as rural.

### Data collection

To investigate effectiveness of the demand optimization intervention, we studied data on laboratory test requests from individual primary medical practices in WHSCT over five consecutive 12 month periods (1 April to 31 March) from 2011-12 (the pre-intervention or ‘baseline’ period), through 2012-2015 (the intervention period), to 2015-16 (the post-intervention period). Test request data were extracted from the laboratory databases of the Altnagelvin Area Hospital (Londonderry), Tyrone County Hospital (Omagh), and the Erne Hospital (subsequently the South West Acute Hospital) (Enniskillen).

The following test groups were studied: 1) profile tests including electrolyte profile, lipid profile, thyroid profile (FT4 and TSH), liver profile, and immunoglobulin profile; 2) glycosylated haemoglobin (HbA_1c_), and 3) prostate-specific antigen (PSA). The number of profile tests (electrolyte profile, lipid profile, thyroid profile, liver profile, immunoglobulin profile) requested in each practice was standardised against the number of registered patients in the practice and expressed as requests per 1000 patients. HbA_1c_ was standardised against the number of patients with diabetes per practice while PSA was standardised against the number of male patients per practice.

Information on individual primary care practices regarding registered patient numbers, the number of male patients, and patients with diabetes was obtained from the Western Health and Social Services Board Integrated Care Partnership system. The patient population served by the 55 practices over the 5-year study period was 316 382 (2011-12), 316 688 (2012-13), 318 057 (2013-14), 319 383 (2014-2015), and 326 429 (2015-2016). The total number of male patients registered in all studied GP practices was 160 046 (2011-12), 152 265 (2012-13), 161 003 (2013-14), 161 824 (2014-2015) and 165 532 (2015-2016) while the number of patients with diabetes was 12 372 (2011-12), 12 871 (2012-13), 13 130 (2013-14), 13 481 (2014-2015) and 14 241 (2015-2016).

Throughout the study period, laboratory tests from primary care were ordered using a paper laboratory request form. All of the tests considered here (with the exception of immunoglobulin profiles) were listed on the request form and were requested by ticking a box on the test request form adjacent to the test name; an immunoglobulin profile was ordered by free text entry on the request form.

### Intervention design

The active intervention was designed to support optimal use of laboratory testing through a quality improvement initiative and took place over the three year period from Apr 2012 to Mar 2015. The intervention package was developed in conjunction with the Western Local Commissioning Group (responsible for commissioning and managing primary care services and consisted of senior primary care doctors). The intervention included several discrete elements. Firstly, awareness of the intervention was promoted through educational sessions on the benefits to patients and clinical teams of the optimal use of laboratory tests. Secondly, educational material was developed in conjunction with primary care clinicians and covered the major clinical indications for a range of most commonly requested tests (i.e. profile tests, HbA1c and PSA) summarised on a single A4 size page. This material together with a document outlining suggested minimum retesting intervals, prepared for the Clinical Practice Group of the Association for Clinical Biochemistry and Laboratory Medicine and supported by the Royal College of Pathologists, were circulated electronically to all GPs. This information was supplemented by face-to-face educational sessions with primary care teams and presentation of data showing the local variability on test requesting rates. Thirdly, all primary care teams were asked to engage in the process of reviewing test requesting procedures within their practice (i.e. what staff are allowed to request tests and what is the process for test requesting), and to reflect on the information provided on their practice test requesting rates and ranking in comparison to other practices. Finally, GPs were further asked to reflect on the appropriateness of their test requesting volume taking into account the educational package, minimum retest intervals and other relevant guidelines.

The Western Local Commissioning Group (WLCG) also made available funding to incentivise participation in the laboratory demand management initiative. All participating primary care practices received a payment of £0.30 per patient registered on their practice list to engage in the process or reviewing and reflecting on test requesting activity. Prior to the intervention each practice received information on its standardised test request rates over the baseline year and its ranking in relation to standardised test request rates of all other practices served by the laboratory.

### Statistical analysis

All statistical analyses were performed using R statistical software, version 3.3.3. Changes in the number of standardised test requests and between-practice variability in standardised test request rates for profile tests, HbA_1c_ and PSA were compared to the pre-intervention (‘baseline’) period (April 2011 – March 2012).

Between-practice variability in ordering of laboratory tests was assessed by calculating the variance (σ^2^). Due to non-normality of the distribution of the standardised number of laboratory test requests caused by the presence of ‘practice outliers’ (i.e. practices with test request rates statistically different from the ordering rates in the other practices), the differences in variances calculated for pre- and post-intervention periods were tested using the Fligner-Killeen (FK) test [29]. The Fligner-Killeen method provides a robust measure, not sensitive to violations of normality, for assessing the homogeneity of variances by ranking the absolute values of differences for each observation from corresponding sample medians [29]. Note that the normality of laboratory test data was tested with the Shapiro-Wilk test [30]. The non-parametric Mann-Whitney-Wilcoxon (MWW) test was used to compare distributions of test request rates from pre- and post-intervention period [31]. In addition, we examined trends in variability of laboratory test ordering using the Mann–Kendall (MK) test [32]. The Mann-Kendall technique is a nonparametric form of monotonic trend regression analysis and hence suitable for not normally distributed data [32]. In all analysis, a *p* < 0.05 was considered significant.

### Governance considerations

This project was undertaken as a quality improvement initiative to promote optimal use of laboratory services. As it was quality improvement initiative rather than a research project, research ethics approval was not considered necessary. WHSCT is the keeper of the laboratory information system data, and all information used within the study was anonymised and not traceable to an individual patient or general practitioner.

## Results

We evaluated the effectiveness of the demand management intervention on the changes in the number and variability in laboratory test request rates for profile tests, HbA_1c_ and PSA by comparing the between-practice differences in test utilization between the pre-intervention (Apr 2011 – Mar 2012) and post-intervention periods (Apr 2015 – Mar 2016) (Fig 1). We found considerable differences across practices and time in test requesting activity.

**Fig 1.**
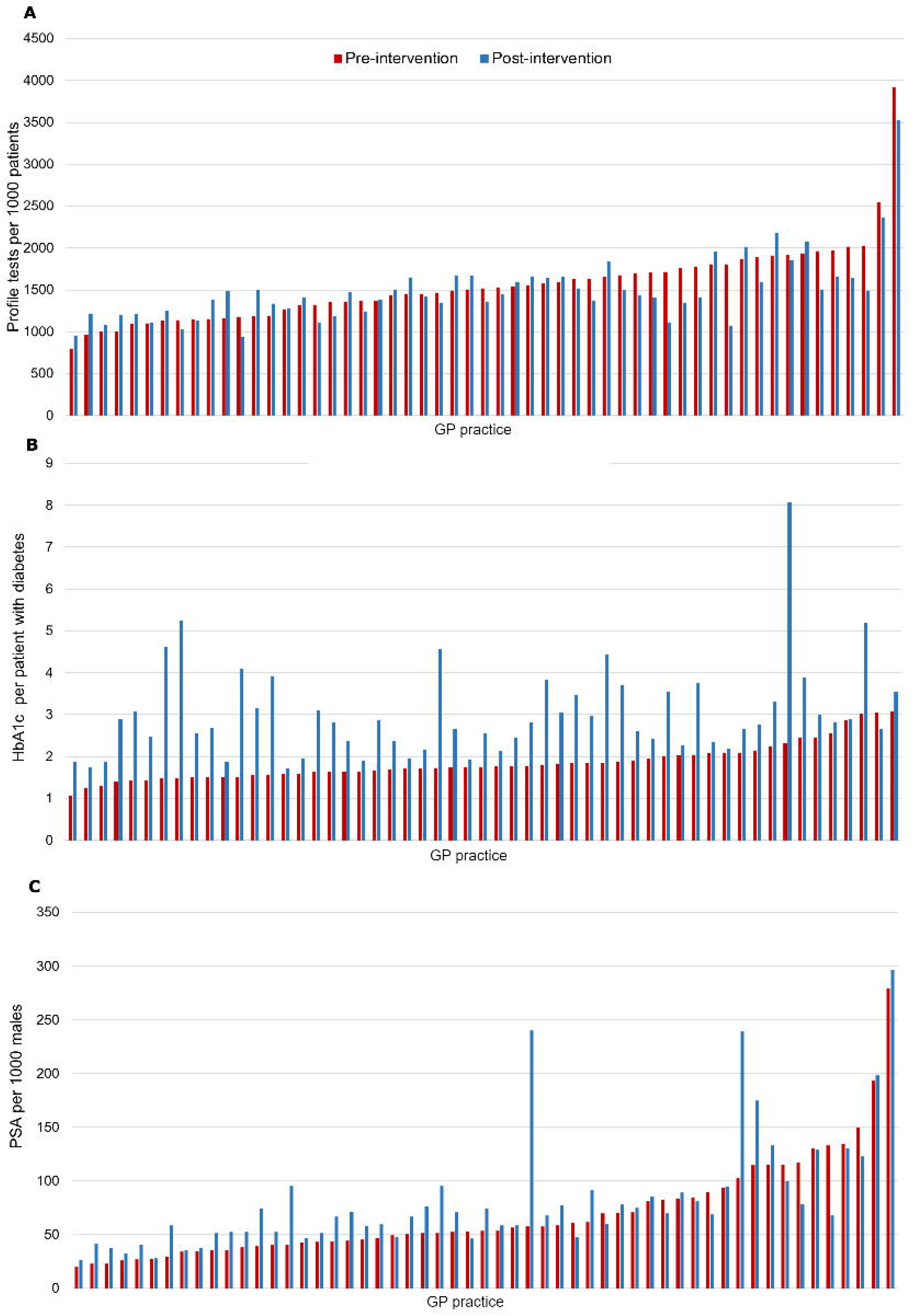
The standardized number of test requests for A) profile test, B) HbA_1c_, and C) PSA in pre-(red) and post-(blue) intervention period for 55 investigated general practices.

### Temporal changes in the standardized number of request rates

The median standardized number of profile test requests for all practices fell from 1519 per 1000 patients pre-intervention to 1441 per 1000 patients one year post intervention (a reduction of 5.1%) (Table 1); however this change was found statistically insignificant (MWW *p* = 0.3) (Table 2). For HbA_1c_, there was a significant increase in the median number of request rates from 1.8 requests per patient with diabetes pre-intervention to 2.8 post-intervention (MWW *p* < 0.0001). The median PSA requests per 1000 male patients increased by 29.3% from 53.2to 68.9 following the intervention (MWW *p* = 0.048) (Table 1 and 2).

**Table 1.** Standardised test request rates for profile tests, HbA_1c_, and PSA over five consecutive 12 month periods (1 April to 31 March) from 2011-12 (the pre-intervention or ‘baseline’ period), through 2012-2015 (the intervention period), to 2015-16 (the post-intervention period.

|  | Pre-intervention |  | Intervention |  | Post-intervention |
| --- | --- | --- | --- | --- | --- |
|  | Apr 2011-Mar 2012 | Apr 2012-Mar 2013 | Apr 2013-Mar 2014 | Apr 2014-Mar 2015 | Apr 2015-Mar 2016 |
| **Profile tests** |  |  |  |  |  |
| Median, (25<sup>th</sup>-75<sup>th</sup> percentile) | 1519, (1230,1765) | 1494, (1280-1762) | 1422, (1247-1658) | 1418, (1277-1606) | 1441, (1244-1644) |
| Mean, (95%CI) | 1554, (1427,1681) | 1556, (1429,1682) | 1499, (1380,1619) | 1485, (1367,1603) | 1498, (1387,1609) |
| (Range) | (798-3919) | (809-4043) | (879-3918) | (868-3840) | (942-3530) |
| Variance | 220152 | 219471 | 195874 | 190917 | 168873 |
| **HbA<sub>1c</sub>** |  |  |  |  |  |
| Median, (25<sup>th</sup>-75<sup>th</sup> percentile) | 1.8, (1.6-2.0) | 2.0, (1.7-2.3) | 2.1, (1.8-2.8) | 2.3, (2.0-2.9) | 2.8, (2.4-3.5) |
| Mean, (95%CI) | 1.9, (1.7,2.0) | 2.0, (1.9,2.2) | 2.3, (2.1,2.5) | 2.6, (2.4,2.9) | 3.0, (2.7 3.3) |
| (Range) | (1.1-3.1) | (1.1-3.4) | (1.3-4.6) | (1.5-6.0) | (1.7-8.1) |
| Variance | 0.2 | 0.3 | 0.6 | 0.8 | 1.2 |
| **PSA** |  |  |  |  |  |
| Median, (25<sup>th</sup>-75<sup>th</sup> percentile) | 53.2, (40.4-84.1) | 59.2, (42.8-101.2) | 59.5, (48.8-89.6) | 63.9, (47.9-83.1) | 68.9, (51.8-90.0) |
| Mean, (95%CI) | 69.4, (56.7,82.0) | 79.2, (62.3,96.0) | 79.6, (60.2,99.0) | 74.8, (62.1,87.4) | 82.9, (68.2,97.6) |
| (Range) | (19.6-279.3) | (19.9-396.1) | (23.1-527.6) | (17.1-274.0) | (26.4-296.9) |
| Variance | 2193 | 3896 | 5154 | 2193 | 2961 |

### Temporal changes in variability of laboratory test ordering

To assess the direction and magnitude of changes in variability of ordering behaviour associated with the intervention, we calculated the variance of test request rates in the post-intervention period and compared it to the pre-intervention data. We also examined the trend in variance across five consecutive time periods, from Apr 2011 – Mar 2012 to Apr 2015 – Mar 2016 (Table 1). The variance for profile test requests fell from σ^2^ = 220152 pre-intervention to σ^2^ = 168873 one year post intervention (a reduction of 23.3%). Despite the fact that this change in variance was found not statistically significant (FK test *p* = 0.2), the Mann–Kendall test indicated the monotonic statistically significant downward trend in variability of profile test request rates (*p* = 0.03) (Table 2). Variance of HbA_1c_ request rates increased from σ^2^ = 0.2 to σ^2^ = 1.2 (an increase of 600%, FK *p* = 0.0001). In addition, we observed a statistically significant upward trend in variance of HbA_1c_ over the study period (MK *p* = 0.03). The between practice variability in the standardized number of PSA requests increased by 35%; however, the reported change was not significant at 95% confidence level (FK *p* = 0.9) (Table 1 and 2).

### Differences in laboratory test requesting between rural and urban practices

Rural practices had a significantly higher median number of profile request rates than urban practices at all time points: baseline, during the intervention and at one year post intervention (Table 3). However, a significant reduction in the median profile test request rates was exclusively observed in rural practices where requests fell by approximately 11.9% (MWW *p* = 0.04) as compared to no significant change in ordering behaviour in urban practices (MWW *p* = 0.9) (Table 2). The median PSA request rates per 1000 male patients increased from 70.4 pre-intervention to 78.2 post-intervention in rural practices (MWW *p* = 0.2) and from 49.2 pre-intervention to 59.4 post-intervention (MWW *p* = 0.1) in urban practices respectively. Given HbA_1c_, we reported a significant increase in the median test requests per patient with diabetes both in rural and urban GP practices; in both cases MWW *p* < 0.0001. It is worth noting that the median of HbA_1c_ request rates was substantially lower that their mean over the whole period of investigation suggesting the presence of practices with ‘outlier’ high request rates.

**Table 2.** Differences in profile test, HbA_1c_, and PSA ordering activity between the pre-(Apr 2011 – Mar 2012) and post-intervention (Apr 2015 – Mar 2016) period. Mann-Whitney-Wilcoxon (MWW) test *p*-value assesses differences between pre-and post-intervention distributions of test request rates. Fligner-Killeen (FK) test *p*-value refers to the significance level of differences in variances. Mann–Kendall (MK) test *p*-value assesses trends in variability of laboratory test ordering. MWW and FK *p*-values < 0.05 indicate significant differences in distribution and variance of test request rates (*). The direction of change in variability of test request rates is indicated by arrows: ↑ for increase and J, for decrease. MK *p*-value < 0.05 implies a monotonic (downward or upward) trend in data (*). MK S-value refers to the direction of the trend i.e. a negative S-value corresponds to the downward trend while a positive S-value indicates an upward trend.

|  | Profile tests | HbA<sub>1c</sub> | PSA |
| --- | --- | --- | --- |
| <u>All</u> |  |  |  |
| Fligner-Killeen test |  |  |  |
| *p*-value | 0.2 ↓ | 0.0001* ↑ | 0.9 ↑ |
| Mann-Whitney-Wilcoxon test |  |  |  |
| *p*-value | 0.3 | < 0.0001* | 0.048* |
| Mann-Kendall test |  |  |  |
| *p*-value | 0.03* | 0.03* | 1 |
| *S*-value | -10 | 10 | 0 |
| <u>Urban</u> |  |  |  |
| Fligner-Killeen test |  |  |  |
| *p*-value | 0.6 ↓ | 0.02* ↑ | 0.9 ↓ |
| Mann-Whitney-Wilcoxon test |  |  |  |
| *p*-value | 0.9 | < 0.0001* | 0.1 |
| Mann-Kendall test |  |  |  |
| *p*-value | 0.2 | 0.2 | 0.5 |
| *S*-value | -6 | 6 | -4 |
| <u>Rural</u> |  |  |  |
| Fligner-Killeen test |  |  |  |
| *p*-value | 0.3 ↓ | 0.008* ↑ | 0.6 ↑ |
| Mann-Whitney-Wilcoxon test |  |  |  |
| *p*-value | 0.04 | < 0.0001* | 0.2 |
| Mann-Kendall test |  |  |  |
| *p*-value | 0.5 | 0.03 | 0.8 |
| S-value | -4 | 10 | 2 |

**Table 3.** A standardised number of profile test, HbA_1c_, and PSA request rates for rural and urban GP practices over five consecutive 12 month periods (1 April to 31 March) from 2011-12 (the pre-intervention or ‘baseline’ period), through 2012-2015 (the intervention period), to 2015-16 (the post-intervention period).

|  | Pre-intervention |  | Intervention |  | Post-intervention |
| --- | --- | --- | --- | --- | --- |
|  | Apr 2011-Mar 2012 | Apr 2012-Mar 2013 | Apr 2013-Mar 2014 | Apr 2014-Mar 2015 | Apr 2015-Mar 2016 |
| **Profile tests** |  |  |  |  |  |
| <u>Rural</u> |  |  |  |  |  |
| Median,<br>(25<sup>th</sup>-75<sup>th</sup><br>percentile) | 1706,<br>(1461-1902) | 1637,<br>(1480-1853) | 1489,<br>(1350-1730) | 1579,<br>(1325-1646) | 1503,<br>(1373-1657) |
| Mean,<br>(95%CI) | 1720,<br>(1486,1953) | 1726,<br>(1493,1960) | 1604,<br>(1370,1837) | 1581,<br>(1350,1813) | 1566,<br>(1359,1774) |
| (Range) | (998-3919) | (1112-4043) | (1139-3918) | (868-3840) | (1073-3530) |
| Variance | 317033 | 314628 | 318440 | 312665 | 251371 |
| <u>Urban</u> |  |  |  |  |  |
| Median,<br>(25<sup>th</sup>-75<sup>th</sup><br>percentile) | 1369,<br>(1168-1642) | 1424,<br>(1212-1644) | 1327,<br>(1215-1596) | 1361,<br>(1211-1530) | 1380,<br>(1211-1619) |
| Mean,<br>(95%CI) | 1426,<br>(1297,1555) | 1424,<br>(1297,1551) | 1418,<br>(1301,1536) | 1410,<br>(1294,1527) | 1444,<br>(1321,1567) |
| (Range) | (798-2543) | (809-2356) | (879-2205) | (893-2297) | (942-2368) |
| Variance | 123789 | 119431 | 102533 | 100589 | 112373 |
| **HbA<sub>1c</sub>** |  |  |  |  |  |
| <u>Rural</u> |  |  |  |  |  |
| Median,<br>(25<sup>th</sup>-75<sup>th</sup><br>percentile) | 1.8,<br>(1.6-2.1) | 2.1,<br>(1.7-2.3) | 2.1,<br>(1.8-2.6) | 2.3,<br>(1.9-2.9) | 2.9,<br>(2.3-3.6) |
| Mean,<br>(95%CI) | 1.9,<br>(1.7,2.1) | 2.1,<br>(1.6,2.3) | 2.3,<br>(2.0,2.7) | 2.8,<br>(2.3,3.3) | 3.2,<br>(2.6,3.8) |
| (Range) | (1.4-3.0) | (1.3-3.1) | (1.4-4.6) | (1.6-6.0) | (1.7-8.1) |
| Variance | 0.2 | 0.3 | 0.7 | 1.4 | 2.2 |
| <u>Urban</u> |  |  |  |  |  |
| Median,<br>(25<sup>th</sup>-75<sup>th</sup><br>percentile) | 1.8,<br>(1.5-1.9) | 1.9,<br>(1.7-2.3) | 2.2,<br>(1.7-2.9) | 2.4,<br>(2.0-2.8) | 2.7,<br>(2.4-3.4) |
| Mean,<br>(95%CI) | 1.8,<br>(1.6,2.0) | 2.0,<br>(1.8,2.2) | 2.3,<br>(2.0,2.6) | 2.5,<br>(2.2,2.7) | 2.9,<br>(2.6,3.1) |
| (Range) | (1.1-3.1) | (1.1-3.4) | (1.3-4.2) | (1.5-3.9) | (1.7-4.6) |
| Variance | 0.2 | 0.3 | 0.6 | 0.4 | 0.5 |
| **PSA** |  |  |  |  |  |
| <u>Rural</u> |  |  |  |  |  |
| Median,<br>(25<sup>th</sup>-75<sup>th</sup><br>percentile) | 70.4,<br>(49.8-115.0) | 70.8,<br>(43.5-124.8) | 72.8,<br>(54.3-107.1) | 74.6,<br>(55.8-102.8) | 78.2,<br>(65.2-105.7) |
| Mean,<br>(95%CI) | 87.4,<br>(62.9,112.0) | 101.9,<br>(67.5,136.3) | 103.1,<br>(60.7,145.4) | 93.4,<br>(68.2,118.7) | 106.8,<br>(77.4,136.2) |
| (Range) | (29.7-279.3) | (35.5-396.1) | (27.8-527.6) | (38.5-274.0) | (35.7-296.9) |
| Variance | 3496.8 | 6880.5 | 10485.7 | 3705.4 | 4997.0 |
| <u>Urban</u> |  |  |  |  |  |
| Median,<br>(25<sup>th</sup>-75<sup>th</sup><br>percentile) | 49.2,<br>(35.5-65.6) | 50.1,<br>(39.7-69.0) | 55.1,<br>(41.0-69.1) | 55.4,<br>(44.3-70.2) | 59.4,<br>(46.5-74.6) |
| Mean,<br>(95%CI) | 55.4,<br>(44.4,66.3) | 61.6,<br>(48.9,74.2) | 61.4,<br>(51.2,71.5) | 60.3,<br>(50.6,70.0) | 64.4,<br>(54.0,74.8) |
| (Range) | (19.6-134.6) | (19.9-170.9) | (23.1-125.8) | (17.1-129.8) | (26.4-132.8) |
| Variance | 891.0 | 1191.4 | 767.6 | 703.8 | 800.5 |

Rural practices had generally higher variance in request rates for profile tests, HbA_1c_, and PSA than urban practices at all time points (Table 2). We did not observe a significant change in variance of profile tests between pre- and post-intervention periods, either in rural (FK *p* = 0.3) or urban (FK *p* = 0.6) practices; however in both cases we reported a downward trend in variance (a σ^2^ reduction of 20.7% and 9.2% respectively). In contrast, a statistically significant change in variance was reported for HbA_1c_ both in rural (FK *p* = 0.008) and urban (FK *p* = 0.02) areas. Despite the 42.9% increase in variance for PSA request rates in rural practices and simultaneous 10.2% decrease in the standardized PSA test requests in urban practices, none of these changes were found statistically significant (Table 2).

## Discussion

While it may be challenging to define an appropriate rate of requesting for most tests, it is certainly difficult to justify very high levels of variability between clinical teams providing care to broadly similar groups of patients within a single healthcare system. This study found high levels of baseline variability between primary care practices in the standardised number of profile tests, HbA_1c_, and PSA, with substantial differences in variability in laboratory test utilization between rural and urban areas. There is little reason to believe that there were significant differences in the characteristics of the practice patient populations within each of rural and urban areas in terms of disease prevalence or morbidity that might explain such high variability. For instance, O’Kane et al. found no link between the number of HbA_1c_ measurements performed per patient with diabetes in practices in an area of N. Ireland and either the reported prevalence of diabetes or Quality and Outcome Frameworks (QOF) scores defining the practice performance in the management of diabetes.

This quality improvement intervention employed to optimise utilization of laboratory tests in primary care was associated with mixed effects. Firstly, there was a reduction of 5.1% in the median profile test requests per 1000 patients (as measured at one year post intervention). This was accounted for entirely by a reduction in rural practices. Secondly, we observed a 23.3% reduction in between practice variability in profile test requesting and this was seen in both urban and rural practices (a decrease in variance of 9.2% and 20.7% respectively). However, during and post-intervention, the standardised profile test request rates and variability continued to be higher in rural than urban practices. Although both the volume and variability in ordering rates for profile tests were reduced, these changes were not statistically significant meaning that the observed differences between the pre- and post-intervention period may have resulted from fluctuation around the baseline or other as yet determined factors. Despite non-significant differences in profile test utilization between the pre- and post-intervention period, the overall statistically significant downward trend in variability (*p* = 0. 03) may indicate a further future decrease in ordering of profile tests.

Given HbA_1c_, we observed a significant increase in both the median number of test requests per patient with diabetes (an increase of 55.6%) as well as in between practice variability (600% increase in variance) between pre- and post-intervention period. Best practice guidelines suggest measuring HbA_1c_ two to three times per year in patients with diabetes and this had been highlighted in the educational material that formed part of the intervention [33]. The increased testing rate may therefore reflect more appropriate monitoring of patients with diabetes. However, as it was not possible to distinguish HbA_1c_ samples which had been requested for diabetes monitoring from those requested for the purposes of diabetes diagnosis, it is difficult to be certain. The use of HbA_1c_ as a diagnostic test for diabetes mellitus had been introduced in 2012 i.e. during the baseline period and it is possible that the increase in requesting reflected its adoption as a diagnostic test rather than as a monitoring test. Yet, the subsequent increase in between practice variability of HbA_1c_ ordering rates may imply that the recommended guidelines on the management of patients with diabetes were not implemented consistently across GP practices.

The increase in the median PSA request rates of 29.3% may be related to the 4.3% rise in the prostate cancer incidence rates in WHSCT over the study period. However, we cannot also exclude the opposite that the increased incidence of prostate cancer may be a consequence of the increased number of PSA requests. In addition, some of the increase in PSA requesting could be associated with more PSA tests being carried out on asymptomatic men, i.e. requests non-compliant with the intervention guidelines. The 35% increase in variability in PSA request rates across general practices between the pre- and post-intervention period could imply greater differential adherence to the national guidelines (e.g. PSA testing not recommended for screening of asymptomatic men) or may have been related to the considerable rise in prostate cancer incidence rates in only some GP practices. However, no evidence data at the individual practice level was available to test such hypothesis.

Although numerous previous studies had documented high degrees of variability in test requesting between primary care teams [34–36], a unique feature of our study was that it assessed the effect of the intervention on the changes in both the volume and between practice variability in test requesting. The relatively poor effectiveness of educational and financial initiatives in diminishing very pronounced differences in test volumes among general practices observed in our study may suggest that the demand optimization intervention undertaken was ineffective. It is clear that the dissemination of clinical management guidelines on appropriate re-testing intervals as well as the benchmarking scheme allowing individual practices to compare their requesting numbers against other practices did not have the effect anticipated. Yet, finding the more suitable interventions may prove to be difficult in the absence of identified factors affecting the variability in test requesting.

A number of initiatives for optimizing demand of laboratory test ordering, aiming at both overutilization and underutilization of tests, have been conducted in primary care. However, the effectiveness of these strategies varied. Several studies reported on mixed effects of educational interventions on laboratory test utilization. Baker et al showed that failure of feedback and educational initiatives to influence the utilization of laboratory tests was associated with baseline performance i.e. how often and how these initiatives were implemented [37]. It is therefore possible that the frequency and form of feedback with guidelines chosen for our study design was not appropriate. In addition, the effect of the demand optimization intervention described here might have been modulated by characteristics of local practices and opinions of individual GPs regarding the role of laboratory tests in patient management. For instance, the observed increase in variability of PSA and HbA_1c_ request rates may indicate that recommended guidelines did not predispose GPs to change their perceptions on the value and role of these tests in patient assessment.

Since the demand optimization intervention showed little effect on laboratory test request rates (e.g. the decrease in variability was only reported for profile tests), other clinical initiatives for optimizing the overall demand and variability of test requests, such as modifications to laboratory requisition forms or introduction of guideline driven decision support systems, should be considered. Previous studies reported on the significant changes in laboratory test ordering behaviour after a redesign of laboratory requisition forms to include fewer test choices [38], and after imposing a clinician-oriented restriction policy on the laboratory test-ordering mechanism (i.e. physicians were required to provide a justification for every test request) [39].

Our study has several limitations worth noting. First, we measured the effect of the demand optimization intervention on changes to laboratory utilization. It is however possible that factors other than the intervention were responsible for utilization changes. Second, since the intervention consisted of several discrete elements (educational sessions, feedback, review of test requesting procedures, financial incentive), it is difficult to determine which element had the largest effect on test requesting rates. Third, we acknowledge that the post-intervention follow-up period might have been too short to determine whether the intervention was in fact ineffective. Finally, increased requesting of laboratory tests does not necessarily translate to decreased appropriateness of their utilization. For instance, the post-intervention increase in median request rates for PSA and HbA_1c_ does not necessarily imply inappropriate testing if it allowed improved patient management. However, the increase in between-practice variability in request rates for those two tests may suggest some degree of inappropriateness in use of laboratory services. Previous work has suggested that large between-practice variability in test utilization is more likely caused by differences in the clinical practice of general practitioners rather than the demographic or socioeconomic characteristics of the practices [36].

Our study has identified considerable variability between general practices in laboratory test request rates and has sought to explore the effect of a demand optimisation intervention on the volume and variability of laboratory test ordering. Future qualitative work could address uncertainty around the factors affecting the variability of test requesting.

## Acknowledgments

The authors would wish to thank the IHAC collaborative network, especially KongFatt Wong-Lin, Colm Hayden, Brendan Bunting and Le Roy Dowey, for helpful discussions; Graham Moore, Austin Tanney, and Paul Barber for computing and technical support; and Stephen Lusty and Peter Devine for administrative support.

## Contributorship

MB performed the analysis and interpretation of the results, and wrote the manuscript. MJO edited the manuscript. MJO, BOH, CM and PC designed and carried out the demand management intervention. MJO wrote the manuscript. BOH, CM and PC edited the manuscript. SA monitored the data collection. All the authors have accepted responsibility for the entire content of this submitted manuscript and approved submission.

